# GoMi - A new gold standard corpus for miRNA Named Entity Recognition to test dictionary, rule-based and machine-learning approaches

**DOI:** 10.1101/2021.10.18.464801

**Authors:** Anika Frericks-Zipper, Markus Stepath, Karin Schork, Katrin Marcus, Michael Turewicz, Martin Eisenacher

## Abstract

Biomarkers have been the focus of research for more than 30 years [REF1]. Paone et al. were among the first scientists to use the term biomarker in the course of a comparative study dealing with breast carcinoma [REF2]. In recent years, in addition to proteins and genes, miRNA or micro RNAs, which play an essential role in gene expression, have gained increased interest as valuable biomarkers. As a result, more and more information on miRNA biomarkers can be extracted via text mining approaches from the increasing amount of scientific literature. In the late 1990s the recognition of specific terms in biomedical texts has become a focus of bioinformatic research to automatically extract knowledge out of the increasing number of publications. For this, amongst other methods, machine learning algorithms are applied. However, the recognition (classification) capability of terms by machine learning or rule based algorithms depends on their correct and reproducible training and development. In the case of machine learning-based algorithms the quality of the available training and test data is crucial. The algorithms have to be tested and trained with curated and trustable data sets, the so-called gold or silver standards. Gold standards are text corpora, which are annotated by expertes, whereby silver standards are curated automatically by other algorithms. Training and calibration of neural networks is based on such corpora. In the literature there are some silver standards with approx. 500,000 tokens [REF3]. Also there are already published gold standards for species, genes, proteins or diseases. However, there is no corpus that has been generated specifically for miRNA. To close this gap, we have generated GoMi, a novel and manually curated <u>**go**</u>ld standard corpus for <u>**m**</u>iRNA. GoMi can be directly used to train ML-methods to calibrate or test different algorithms based on the rule-based approach or dictionary-based approach. The GoMi gold standard corpus was created using publicly available PubMed abstracts.

GoMi can be downloaded here: https://github.com/mpc-bioinformatics/mirnaGS---GoMi.

## 1. Introduction

### 1.1 miRNA as research target

Micro RNA (miRNA) are short non-coding RNA fragments that have usually 21-23 nucleotides. miRNAs do not encode proteins but are involved in gene expression via transcriptional or translational pathways. [REF4] MiRNA was first observed in C.Elegans in the late 20th century.[REF6] In the course of protein translation, miRNA can inhibit various substances or degrade messenger RNA (mRNA). Their biogenesis is influenced by different factors such as mutations or phosphorylation. These factors are also influenced by environmental factors such as stress or certain diseases.[REF5] Therefore, miRNAs can also function as biomarkers to diagnose diseases, to follow the course of a disease or to evaluate the success of a therapy. A well researched example is breast cancer. The expression values of different miRNAs can provide information about the stage of the disease and serve as a prognostic marker, while miRNA-200a can also be used for diagnosis. [RE6] In recent years, research projects have focused on miRNA, in particular regarding the relationship between viral diseases and miRNA, e.g. for COVID-19. In the context of a SARS-COV-2 infection, miRNAs were found to be potential biomarkers [REF2] and were also investigated in the course of the vaccine developed by the company **BioNTech SE**. [REF4]. In addition to SARS-Cov-2 infections, other viral diseases are also associated with miRNA exosomes, such as influenza A.[REF7]. The increasing interest in miRNA research is reflected by the related publication activity. In PubMed there are 125,030 publications (as of July 2021) tagged with the keywords “miRNA” or “micro RNA”. Using biomedical text mining, additional information and/or knowledge may be automatically extracted from this literature.

### 1.2 Natural Language Processing in biomedical texts

Natural Language Processing (NLP) includes different algorithms which are all related to text processing, including the manipulation of texts or the extraction of certain keywords from a text. These could be from medical texts but also from any other subject area. [REF9]. Two well-known aspects in NLP are Named Entity Recognition (NER) and Relation Extraction (RE). The first refers to the process of identifying and tagging a word or groups of words of interest in a given text with the correct entity class. For example, one task may be to tag all surface proteins in a scientific article. RE is used to identify the relationship between the tagged entities. [REF9] An example for RE is BIONDA, a biomarker database that extracts the relations between diseases and their potential biomarkers based on a sentence-wise co-occurrence approach. [REF10] However, obviously a correct NER is a crucial prerequisite for the RE.

The different approaches for NER can be divided into three categories, dictionary-based approaches, rule-based approaches and machine learning-based approaches.

The dictionary-based approach uses dictionaries, i.e. lists of terms, which are searched in the texts. The texts are examined word by word and it is checked whether they occur in the dictionary. This approach is usually very time-consuming due to the brute force approach and achieves worse recall scores than other approaches due to its strictness. A fuzzy search can help to make the approach more flexible but can also lead to wrong results being found. However, many NER models, especially older ones, use the dictionary-based approach because of well annotated dictionaries in the biomedical domain and because of their simplicity and availability [REF 11]. E.g. high-quality sources for dictionaries for diseases, the Disease Ontology, for genes/proteins UniProt and for miRNA mirBase are available. [REF12,13,14]. Rule-based approaches use self-defined rules to tag and recognize words.These rules are based on different patterns. The creation of these rules requires expert knowledge and there is a risk that the rules are kept too simple and the algorithm loses specificity due to this. On the other hand, the rules can also be too strict and therefore lead to poor recall [REF7,8]. A successful implementation of a rule-based approach is offered by DrNER [REF15], a model developed by Eftimov et al. to extract dietary recommendations from unstructured texts. The rules are based on chemical notations and regular expressions. The construction of such rules was very time consuming. However, the results are satisfactory and the authors reported a precision of 99% and a recall of 96% for the category “FOOD” from scientifically validated websites, scientific publications and other text corpora [REF15].

Finally, as part of the machine learning-based approach, trained neural networks recognize the words of interest and tag them. Different concepts and network architectures have been published. REF16] While Google uses a transformer architecture with BERT, HunFlair uses an NLP framework[REF17]. HunFlair is a software library that offers various algorithms. On the one hand it is based on Huner approach with a pre-trained languages module. On the other hand, it contains the framework Flair which includes methods for labelling, training and other classifications for text sequences.[REF17] As transformer networks PubMedBert and BioBert are offered for the biomedical area, where these differ only in the supplied training data for pre-training, while BioBert uses Wikipedia books as training data, PubMedBert uses exclusively PubMed abstracts.[REF16]

### 1.3 Silver and gold standards for text mining

Training data is enormously important to achieve good NER results with machine-learning approaches. Therefore, it is important to use correctly annotated training data.[REF18] In the literature, a distinction is made between gold and silver standards. Silver standards are automatically annotated by algorithms and are therefore not placed on the same level as gold standards, which are manually curated by experts. For the silver standard for phenotype recognition published by Oellrich et al. an F-Score between 0.5 and 0.6 was observed outperforming other three competitor tools in almost all cases. [REF19]

On the other hand, gold standards are more reliable due to the manual curation by experts and are therefore used to validate NER approaches or to train neural networks. In the literature, there are already some NER gold standard datasets like NCBI-Disease for diseases [REF20] or JNLPBA for genes/proteins [REF21]. The number of tokens ranges from about 80,000 to 3,700 for the gold standards presented by Habibi et al. BCSCHEMD is the largest with 79,852 tokens and deals with chemicals and the smallest dataset is Species-800 with 3,708. Three of nine datasets described in [REF22] deal with diseases, the others with species, drugs or genes/proteins. [REF22]. In addition, there are already gold standards that deal with relation extraction or question answering, but these are relatively small corpora ranging between 355 and 10,035 tokens [REF23]. In total there are many different gold standards for various questions. But there is no known dataset that has been specifically generated for miRNAs. Hence, there is a need for such a corpus, e.g., to train a machine learning-based NER-model tagging only miRNAs in a given text. Moreover, also miRNA-specific rule-based and dictionary-based approaches could be assessed and compared using such a corpus. Among other things, GoMi also contains a development dataset which is required for validation in the ML approaches.

## 2. Methods

### 2.1 Structure of the dataset

GoMi contains 169,995 tokens. These are divided into subsets for testing and training of NER algorithms. For the neuronal network use case, ten successive training datasets have been formed out of the training part of GoMi. The 10th and therefore largest training dataset contains all the others. The test part of GoMi has also been split into ten non-overlapping test datasets. They are also stored as a complete version under Final_Test_Set on GitHub. This structure makes it possible for the user to combine the different test and train splits and therefore to adapt GoMi to their individual questions and data.

### 2.2 Information retrieval and text preprocessing

All article abstracts annotated here were downloaded from PubMed [REF24] in the PubMed text format. Since the novel corpus was also aimed to evaluate the biomarker database BIONDA, the query to retrieve the article abstracts was set to “mirna biomarker”. This ensured that only abstracts relevant to BIONDA were used to generate the corpus. The abstracts were downloaded via the web interface of PubMed.

In order to annotate the abstracts so that they can be used later for validation or the training of ML-models, they must be tokenized. This means that the abstract is taken apart word by word. The desired output is then a.tsv file in which each word contained in the abstract is placed in a single row. To accomplish this task the Stanza package[REF25] was used and integrated in a Python script implemented for this purpose.

### 2.3 Manual annotation of PubMed abstracts

After tokenization of the abstracts, they were annotated manually. The annotation is done according to the CoNLL-U annotation format [REF26] in combination with the BIO principle [REF21]. The CoNLL-U format is a way of structuring texts, which was defined at the Conference on Computational Natural Language Learning (CoNLL). The annotations are encoded in UTF-8 format and stored as plain text without further formatting. The original format consists of 10 columns: ID,FORM, LEMMA, UPOS, XPOS, FEATS,HEAD,DEPREL, DEPS and MISC. For GoMi, a modification was used and only the LEMMA, i.e. the actual word, was used [REF26] The gold standards already mentioned above, such as NCBI-Disease or JNLPBA, also were annotated following the BIO annotation format [REF 20,21]. Here, a “B” marks the beginning of an entity, an “I” stands for words inside an entity and an “O” represents words outside entities. To demonstrate this, table 1 shows an annotated example from the novel miRNA corpus. In order to ensure the correctness of the manual annotation, it was performed twice. Finally, the annotated GoMi dataset was structured as follows: there are 10 different training sub-corpora and 10 different test sub-corpora in different sizes, which have been predefined.The users can therefore assemble their set from the total data in such a way that they resemble their data to calibrate a network or to test a NER algorithm.

**Table 1:**
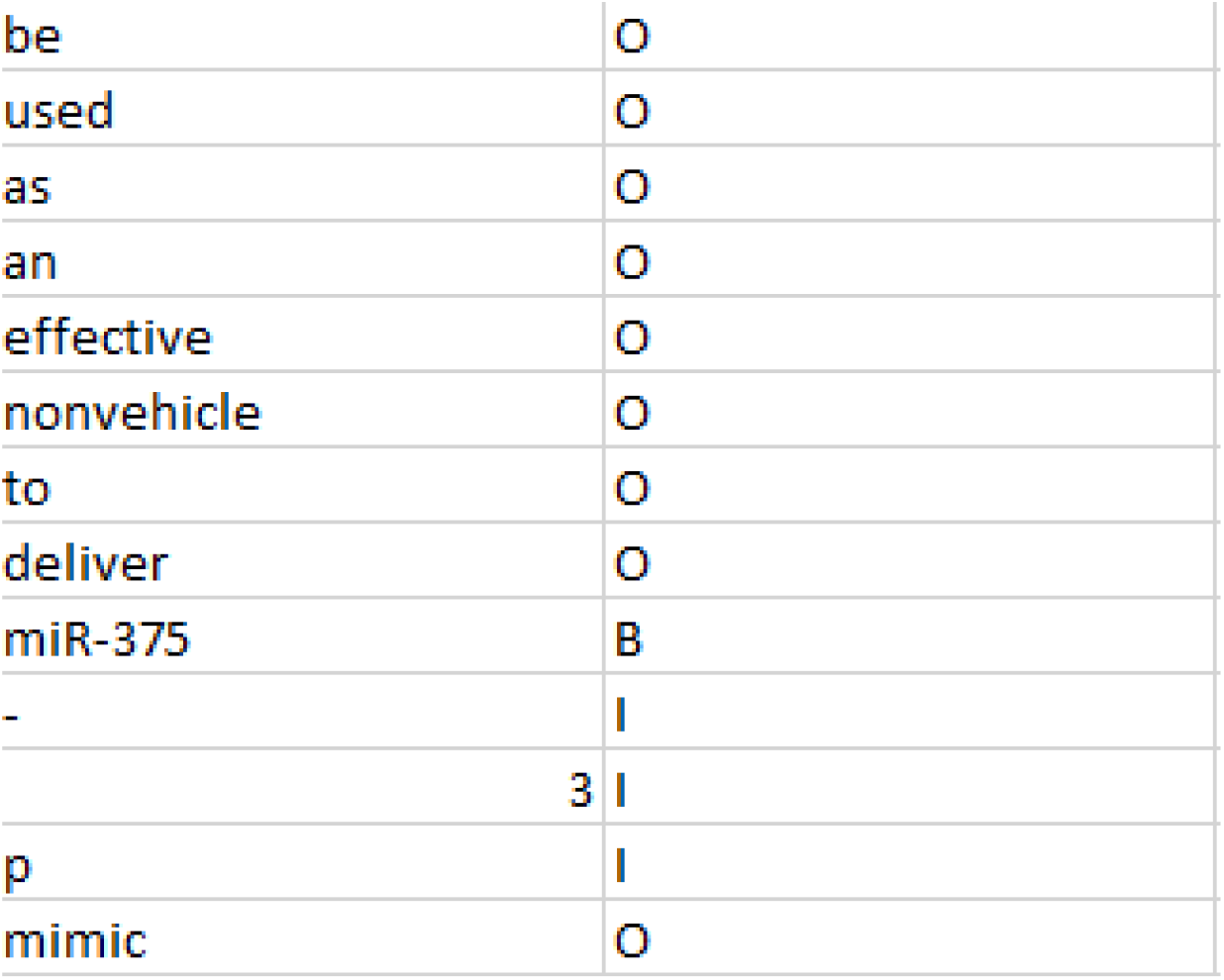
Example from the first test dataset. In the left column the single words of an abstract obtained from the tokenizer are listed. In the right column the annotation following the BIO format and performed by the curator is shown, where a single miRNA entity was tagged. Here, the “B” marks the beginning of the miRNA entity, “I” is used for words that are inside the entity and “O” represents all tokens that are outside the miRNA entity.

### 2.4 Training of the data set in the neural network

All 10 training data and test sub-corpora of GoMi, as well as the devel dataset, have been used to re-train and evaluate the resulting PubMedBERT and BioBERT network models. This was implemented in Python in order to test the usability of GoMi and whether it may be used to improve the training of neural network models that are focused on miRNA. For both network training procedures a 10-fold cross-validation was used. With this, recall, precision and the F-Score were calculated [REF27] Finally, the runtime of both networks was recorded and compared.

### 2.5 Hardware-Setup

The networks were performed in Google Colab with the setup shown in table 2.

**Table 2:**
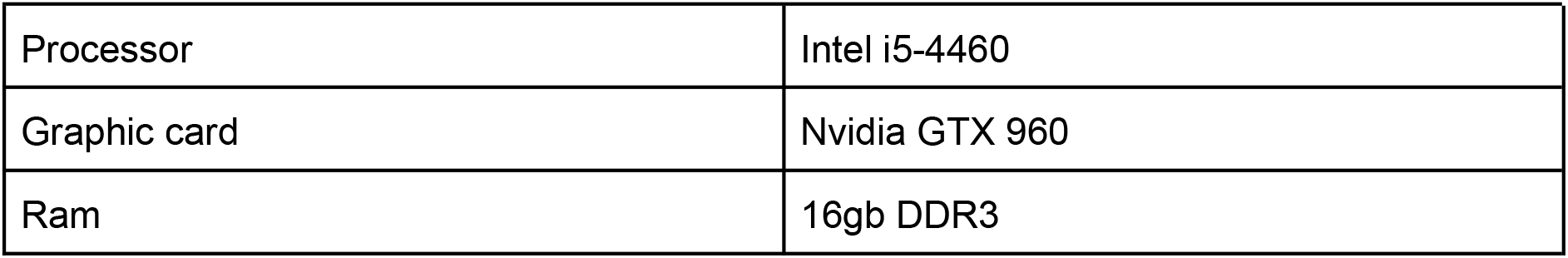
PC setup for the neuronal networks

## 3. Results

In order to generate GoMi, in total 600 PubMed abstracts with approx. 49,000 sentences have been downloaded and annotated. As shown in table 2, this results in different token numbers for the individual training and test sub-corpora. In order to test the data set in different combinations, it was divided into 10 parts as shown in table 2. For later use, it is up to the users to combine these sub-corpora according to their requirements. The records for the training data range from 145,373 to 10,974 tokens.

To test the novel GoMi corpus as a use case for the training of neural network-based NER models, it was split up as explained in 2.1. The number of tokens increases in steps of approx. 10,000 tokens from sub-corpus to sub-corpus. The test sub-corpora is completely distinct from the training sub-corpora. On the other side, the training sub-corpora build on each other, i.e. the previous dataset is always part of the next one. This applies to all 10 sets, therefore the 10th training sub-corpus contains the union of all previous GoMi training subsets. The test sub-corpora were constructed differently, here we decided that all of them should be independent, due to their smaller size. The sum represents the complete GoMi corpus, which can be found as “Complete_Test” in Github.

The training data sub-corpora contain 628 and 2,702 and the test sub-corpora contain 8 to 229 annotations. For our test purposes, the corpus was divided as shown in table 3 and 10 different combinations were tested. This is a combination of the 5th test dataset and all existing training datasets. The supplement contains an Excel sheet with the calculations for the test runs of the re-trained PubMedBERT and BioBERT network models. In addition to the F-Score, recall, precision, runtime and loss are also recorded. For PubMedBert, recall and precision range from 0.94 to 0.7 for all runs except for the run with the smallest training data set, where precision and recall values of 0.41 and 0.53 were achieved. BioBert, on the other hand, achieved precision and recall values between 0.8 and 0.9 for every run except for run 6. In the sixth run, only a precession of 0.625 was achieved. GoMi thus achieves similarly good values as in the BioCreative Study, although no miRNA dataset was tested here.

**Table 3:** Overview of all annotated tokens and tokens in general of the training and test sub-corpora of GoMi.

|  | Tokens |  | Annotations |  |
| --- | --- | --- | --- | --- |
|  | Train | Test | Train | Test |
| 1 | 10,974 | 569 | 628 | 8 |
| 2 | 21,216 | 1,174 | 1,236 | 23 |
| 3 | 43,717 | 3,499 | 2,024 | 96 |
| 4 | 86,032 | 7,293 | 2,106 | 59 |
| 5 | 96,594 | 9,789 | 2,208 | 125 |
| 6 | 108,221 | 13,260 | 2,326 | 110 |
| 7 | 118,671 | 18,189 | 2,441 | 129 |
| 8 | 129,806 | 20,299 | 2,566 | 149 |
| 9 | 137,183 | 22,314 | 2,603 | 277 |
| 10 | 145,373 | 24,622 | 2,702 | 228 |
| Sum | 145,373 | 100,928 | 2,702 | 1,204 |

Figure 1 shows the F-score of 10 runs for the combination of the 5th test dataset against all available training data. The 5th test dataset contains 9,789 tokens including 125 annotated miRNA tokens.

**Figure 1:**
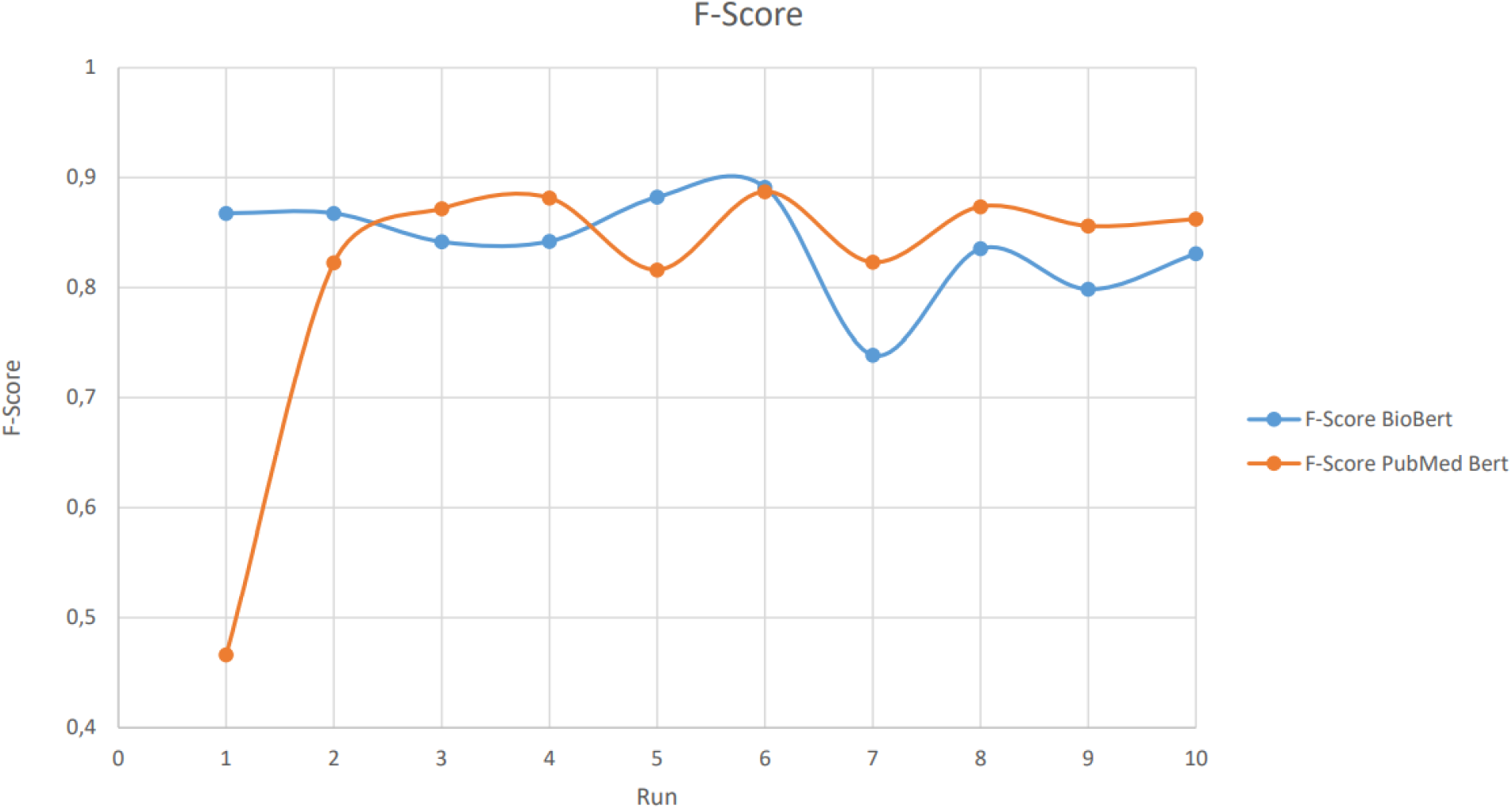
The graph shows the F-score calculated by PubMedBert and BioBert for 10 runs of the re-trained network models using the 5th test sub-corpus against all training sub-corpora.

The highest F-score is achieved by PubMedBERT and BioBERT with the 6th training sub-corpus. For the small training sub-corpora 1 and 2, PubMedBERT shows a poorer performance than BioBERT, which ranges 0.4 - 0.8 than BioBERT. On the other hand, BioBERT is approx. 0.1 point worse on the larger training sub-corpora and performs 0.73-0.83 in contrast to PubmedBERT with 0.82-0.87.

In general, the re-training runtime of BioBert was always lower than that of PubMedBERT (Fig.2). In runs 4-8 both have an identical runtime of 13 minutes, which is also maintained in the last runs of BioBERT. PubMedBert, on the other hand, achieves a runtime of 14 minutes in runs 9 and 10.

**Figure 2:**
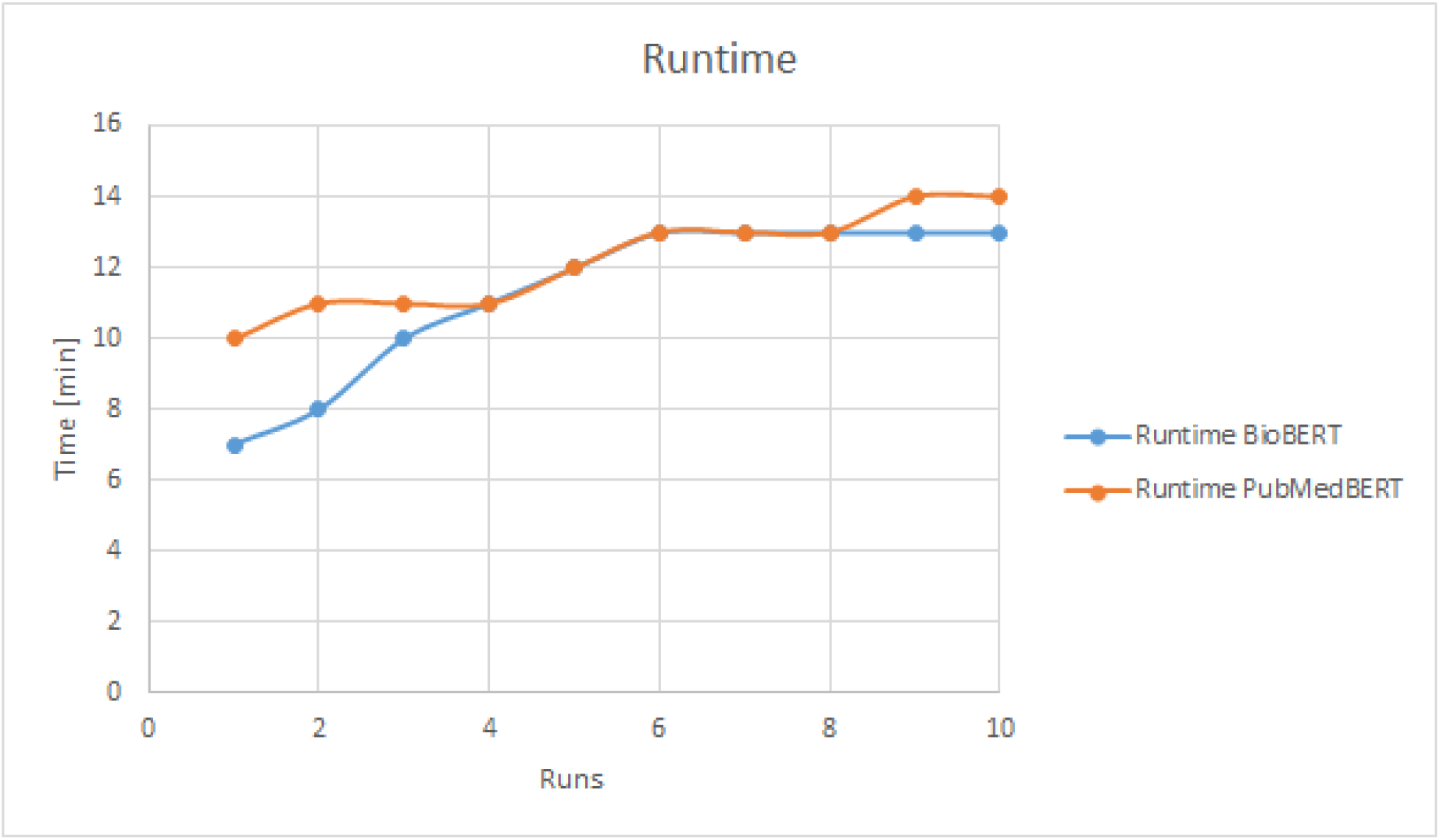
Re-training runtime for PubMedBERT and BioBERT

## 4. Discussion

The GoMi corpus presented here contains 139,885 tokens in total, of which 22,542 are annotated as miRNAs. Hence, this expert curated corpus represents a medium sized dataset compared to other datasets in the biomedical field as can be seen in Habibi et al [REF22]. However, it is much larger than the Species-800 [REF28] and the LINNAEUS[29] data sets. However, it is smaller than the commonly used corpora for chemicals like BC4CHEMD [REF30], which also include many other entity classes than miRNA. On the other hand, the number of annotated miRNA corresponds to corpora for proteins/genes, such as JNLPBA with 35,460 annotations and BC2GM with 20,703 annotations [REF30,31]. However, in these two datasets the focus is much more narrow as proteins and genes are tagged and thus they are not useful for testing an approach that is completely specialized to miRNA. To our knowledge, there is no comparable gold standard dataset like GoMi, which focuses explicitly on miRNA. GoMi offers valuable data for various NLP tasks in the biomedical field as a stand alone dataset or in combination with other gold standards like BC4CHEMD. Due to the original intention to use GoMi for testing the biomarker database BIONDA only PubMed abstracts were used since BIONDA contains only information extracted from these texts; an extension to this would be the annotation of clinical patient data, as in the publication Akhondi et. al [**REF29**]. Another extension could be the annotation of preprints or full text articles.

Furthermore, the datasets in the PubMedBert and BIoBert networks achieve comparable F-scores as presented in the BioCreative Study [REF34]. PubMedBERT achieves higher values, which could be due to the fact that the annotated articles are also PubMed articles and PubMedBERT was finally trained on exactly this data type. With smaller training data, however, BioBERT is better, which could be related to the fact that BioBERT has generally been trained with more training data than PubMedBERT.

As a conclusion, GoMi is a large and well annotated gold standard corpus to calibrate the setting in neural networks and to test dictionary, rule-based and machine learning approaches.

## Supporting information

Supplemental Calucation BioBert Neuronal Network

Supplemental Calucation PubmedBert Neuronal Network

## Data availability

GoMi can be downloaded here: https://github.com/mpc-bioinformatics/mirnaGS---GoMi.

## Acknowledgement

This work was supported by the German Network for Bioinformatics Infrastructure (de.NBI), a project of the German Federal Ministry of Education and Research (BMBF) [FKZ 031 A 534A]. The funding of M.E. relates to PURE and VALIBIO, projects of Northrhine-Westphalia.

