## Supplemental Calucation BioBert Neuronal Network for "GoMi - A new gold standard corpus for miRNA Named Entity Recognition to test dictionary, rule-based and machine-learning approaches"

| | | | | | | |
|---|---|---|---|---|---|---|
| Datensatz | Token | Annotation | Eval | loes | | Precission |
| 1 | 11061 | 138 | | | 0,0268958672 | 0,8260869565 |
| 2 | 21303 | 608 | | | 0,03789 | 0,8024691 |
| 3 | 32830 | 1396 | | | 0,0370688656 | 0,8214285714 |
| 4 | 42955 | 1478 | | | 0,0321453218 | 0,83125 |
| 5 | 53517 | 1580 | | | 0,0423953279 | 0,813253012 |
| 6 | 65145 | 1698 | | | 0,034499415 | 0,8209876543 |
| 7 | 75595 | 1813 | | | 0,0658823816 | 0,625 |
| 8 | 86730 | 2566 | | | 0,0293281813 | 0,8602941176 |
| 9 | 94107 | 1955 | | | 0,066350513 | 0,7348066298 |
| 10 | 102297 | 2037 | | | 0,0525553237 | 0,8011695906 |
| | | 628 | | | | |
| | | | | | 10242 | 11527 |
| | Testdaten | | Train | | | |
| | Token | Annotation | Token | Annotation | | Eval |
| 1 | 569 | 8 | 145373 | | 2702 | |
| 2 | 1174 | 23 | 145373 | | 2702 | |
| 3 | 3499 | 96 | 145373 | | 2702 | |
| 4 | 7293 | 59 | 145373 | | 2702 | |
| 5 | 9789 | 125 | 145373 | | 2702 | normaltestset |
| 6 | 13260 | 110 | 145373 | | 2702 | |
| 7 | 18189 | 129 | 145373 | | 2702 | |
| 8 | 20299 | 149 | 145373 | | 2702 | |
| 9 | 22314 | 277 | 145373 | | 2702 | |
| 10 | 24622 | 228 | 145373 | | 2702 | |

| | | | | | | |
|---|---|---|---|---|---|---|
| COVID | All | Disease | mirna | loss | | Precission |
| train | 452 | 314 | 138 | | 0,0533927238 | 0,8554216867 |
| test | 56 | 6 | 10 | | | |
| devel | 157 | 121 | 36 | | | |

Biobert

| Recall | F-Score | Test | TIME |
|---|---|---|---|
| 0,8636363636 | 0,8444444444 | | 7 |
| 0,84415844 | 0,8227848 | | 8 |
| 0,8961038961 | 0,8571428571 | | 10 |
| 0,8636363636 | 0,847133758 | | 11 |
| 0,8766233766 | 0,84375 | | 12 |
| 0,8636363636 | 0,8417721519 | | 13 |
| 0,8527131783 | 0,7213114754 | | 13 |
| 0,9069767442 | 0,8830188679 | | 13 |
| 0,8636363636 | 0,7940298507 | | 13 |
| 0,8896103896 | 0,8430769231 | | 13 |
| 10125 | 10562 | 11628 | 10450 |

| loss | Precission | Recall | F-Score |
|---|---|---|---|
| 0,0596537243 | 0,7784090909 | 0,8896103896 | 0,8303030303 |
| 0,0619925776 | 0,7555555556 | 0,8831168831 | 0,8143712575 |
| 0,0497840621 | 0,8081395349 | 0,9025974026 | 0,8527607362 |
| 0,0599904794 | 0,7941176471 | 0,8766233766 | 0,8333333333 |
| 0,0525553237 | 0,8011695906 | 0,8896103896 | 0,8430769231 |
| 0,0559148495 | 0,7885714286 | 0,8961038961 | 0,8961038961 |
| 0,0546305858 | 0,816091954 | 0,9220779221 | 0,8658536585 |
| 0,0586787579 | 0,8068181818 | 0,9220779221 | 0,8606060606 |
| 0,0572550838 | 0,8323699422 | 0,9350649351 | 0,880733945 |
| 0,0566967961 | 0,816091954 | 0,9220779221 | 0,8658536585 |

| Recall | F-Score | Test | TIME |
|---|---|---|---|
| 0,8819875776 | 0,8685015291 | | |

Biobert

| loes | Precission | Recall | F-Score |
|---|---|---|---|
| 0,037899387 | 0,8251748252 | 0,9147286822 | 0,8676470588 |
| 0,04232099 | 0,8251748251 | 0,914728 | 0,86764 |
| 0,0503977909 | 0,7852348993 | 0,9069767442 | 0,8417266187 |
| 0,033877497 | 0,8175182482 | 0,8682170543 | 0,8421052632 |
| 0,0389095788 | 0,8391608392 | 0,9302325581 | 0,8823529412 |
| 0,0373116602 | 0,8367346939 | 0,9534883721 | 0,8913043478 |
| 0,0794688574 | 0,6615384615 | 0,8376623377 | 0,7392550143 |
| 0,0545499216 | 0,8148148148 | 0,8571428571 | 0,835443038 |
| 0,039651167 | 0,7569444444 | 0,8449612403 | 0,7985347985 |
| 0,0375093932 | 0,7902097902 | 0,8759689922 | 0,8308823529 |
| 11135 | 7377 | | |

| Test | TIME | loss | Precission |
|---|---|---|---|
| | | 0,0693446212 | 0,5454545455 |
| | | 0,1471314682 | 0,547826087 |
| | | 0,1471314682 | 0,547826087 |
| | | 0,0219884118 | 0,8125 |
| | | 0,0375093932 | 0,7902097902 |
| | | 0,0179082698 | 0,8650793651 |
| | | 0,0550799495 | 0,7516339869 |
| | | 0,0294415539 | 0,8253012048 |
| | | 0,0206077115 | 0,8647798742 |
| | | 0,0177008533 | 0,844 |

| loss | Precission | Recall | F-Score |
|---|---|---|---|
| 0,0384305718 | 0,9423076923 | 0,8448275862 | 0,8909090909 |

Biobert

| Testset | Token | Annotation | Devel | Token | Annotation |
|---|---|---|---|---|---|
| | 9789 | 125 | | 10974 | 629 |

| Recall | F-Score |
|---|---|
| 0,75 | 0,631578947 |
| 0,6428571429 | 0,591549296 |
| 0,6428571429 | 0,591549296 |
| 0,8813559322 | 0,845528455 |
| 0,8759689922 | 0,830882353 |
| 0,9646017699 | 0,912133891 |
| 0,8778625954 | 0,809859155 |
| 0,8954248366 | 0,858934169 |
| 0,9649122807 | 0,912106136 |
| 0,9213973799 | 0,881002088 |
