## Supplemental Calucation PubmedBert Neuronal Network for "GoMi - A new gold standard corpus for miRNA Named Entity Recognition to test dictionary, rule-based and machine-learning approaches"

| Datensatz | Token | Annotation | Eval | loss |
|---|---|---|---|---|
| 1 | 10974 | 628 | | 0,0425365985 |
| 2 | 21216 | 1236 | | 0,04232009 |
| 3 | 43717 | 2024 | | 0,0357046809 |
| 4 | 86032 | 2106 | | 0,0358501764 |
| 5 | 96594 | 2208 | | 0,0379079216 |
| 6 | 108221 | 2326 | | 0,047029392 |
| 7 | 118671 | 2441 | | 0,0487581262 |
| 8 | 129806 | 2566 | | 0,0605768218 |
| 9 | 137183 | 2603 | | 0,0542484468 |
| 10 | 145373 | 2702 | | 0,0531034749 |

| | Testdaten | | Train | | |
|---|---|---|---|---|---|
| | Token | Annotation | Token | Annotation | Eval |
| 1 | 569 | 8 | 145373 | 2702 | |
| 2 | 1174 | 23 | 145373 | 2702 | |
| 3 | 3499 | 96 | 145373 | 2702 | |
| 4 | 7293 | 59 | 145373 | 2702 | |
| 5 | 9789 | 125 | 145373 | 2702 | normaltestset |
| 6 | 13260 | 110 | 145373 | 2702 | |
| 7 | 18189 | 129 | 145373 | 2702 | |
| 8 | 20299 | 149 | 145373 | 2702 | |
| 9 | 22314 | 277 | 145373 | 2702 | |
| 10 | 24622 | 228 | 145373 | 2702 | |

| COVID | All | Disease | mirna | loss | Precission |
|---|---|---|---|---|---|
| train | 452 | 314 | 138 | 0,0533927238 | 0,8554216867 |
| test | 56 | 46 | 10 | | |
| devel | 157 | 121 | 36 | | |

PubmedBert

| Precission | Recall | F-Score | Test | TIME |
|---|---|---|---|---|
| 0,6524390244 | 0,6948051948 | 0,6729559748 | | 10 |
| 0,8251748 | 0,91472862 | 0,867647 | | 11 |
| 0,8322981366 | 0,8701298701 | 0,8507936508 | | 11 |
| 0,8481012658 | 0,8701298701 | 0,858974359 | | 11 |
| 0,81875 | 0,8506493506 | 0,8343949045 | | 12 |
| 0,8303030303 | 0,8896103896 | 0,8589341693 | | 13 |
| 0,8083832335 | 0,8766233766 | 0,8411214953 | | 13 |
| 0,7906976744 | 0,8831168831 | 0,8343558282 | | 13 |
| 0,8106508876 | 0,8896103896 | 0,8482972136 | | 14 |
| 0,8303030303 | 0,8896103896 | 0,8589341693 | | 14 |

| loss | Precission | Recall | F-Score | Test |
|---|---|---|---|---|
| 0,0597482017 | 0,8414634146 | 0,8961038961 | 0,8679245283 | |
| 0,0503381546 | 0,8343195266 | 0,9155844156 | 0,8730650155 | |
| 0,0515973891 | 0,8117647059 | 0,8961038961 | 0,8518518519 | |
| 0,0503201484 | 0,8502994012 | 0,9220779221 | 0,8847352025 | |
| 0,0531034749 | 0,8303030303 | 0,8896103896 | 0,8589341693 | |
| 0,0512129731 | 0,8197674419 | 0,9155844156 | 0,8650306748 | |
| 0,059979784 | 0,8275862069 | 0,9350649351 | 0,8780487805 | |
| 0,0584445791 | 0,8103448276 | 0,9155844156 | 0,8597560976 | |
| 0,0605917832 | 0,8181818182 | 0,9350649351 | 0,8727272727 | |
| 0,0606839494 | 0,8313953488 | 0,9285714286 | 0,8773006135 | |

| Recall | F-Score | Test | TIME | loss |
|---|---|---|---|---|
| 0,8819875776 | 0,8685015291 | | | 0,038430572 |

### PubmedBert

| loss | Precission | Recall | F-Score |
| --- | --- | --- | --- |
| 0,0647756721 | 0,4131736527 | 0,5348837209 | 0,4662162162 |
| 0,0378291724 | 0,8042469 | 0,844155844 | 0,82278548 |
| 0,0483788033 | 0,8263888889 | 0,9224806202 | 0,8717948718 |
| 0,041242683 | 0,8439716312 | 0,9224806202 | 0,8814814815 |
| 0,0475882388 | 0,7762237762 | 0,8604651163 | 0,8161764706 |
| 0,047029392 | 0,8356164384 | 0,9457364341 | 0,8872727273 |
| 0,0387347421 | 0,7702702703 | 0,8837209302 | 0,8231046931 |
| 0,0457707007 | 0,8175675676 | 0,9379844961 | 0,8736462094 |
| 0,0405145101 | 0,8169014085 | 0,8992248062 | 0,8560885609 |
| 0,0410645972 | 0,8095238095 | 0,9224806202 | 0,8623188406 |

| TIME | loss | Precission | Recall |
| --- | --- | --- | --- |
| | 0,1850295663 | 0,625 | 0,625 |
| | 0,2743710064 | 0,4 | 0,2608695652 |
| | 0,1372613364 | 0,58 | 0,5918367347 |
| | 0,0158909501 | 0,875 | 0,9491525424 |
| | 0,0410645972 | 0,8095238095 | 0,9224806202 |
| | 0,0142152314 | 0,9159663866 | 0,9646017699 |
| | 0,0671562825 | 0,7435897436 | 0,8854961832 |
| | 0,0322136294 | 0,8154761905 | 0,8954248366 |
| | 0,0232434916 | 0,8690095847 | 0,9543859649 |
| | 0,0195664503 | 0,83203125 | 0,9301310044 |

| Precission | Recall | F-Score |
| --- | --- | --- |
| 0,9423076923 | 0,8448275862 | 0,8909090909 |

PubmedBert

| Testset | Token | Annotation |
|---|---|---|
| | 9789 | 125 |

F-Score

| F-Score |
|---|
| 0,625 |
| 0,3157894737 |
| 0,5858585859 |
| 0,9105691057 |
| 0,8623188406 |
| 0,9396551724 |
| 0,8083623693 |
| 0,8535825545 |
| 0,9096989967 |
| 0,8783505155 |

| Devel | Token | Annotation |
| --- | --- | --- |
| | 10974 | 629 |
